# Sequential enrichment at the nuclear periphery of H2A.Zac and H3K9me2 accompanies pluripotency loss in human embryonic stem cells

**DOI:** 10.1101/2020.02.15.951103

**Authors:** Georgia Rose Kafer, Regina Rillo-Bohn, Peter M. Carlton

**Affiliations:** Institute for Integrated Cell-Material Sciences (iCeMS), Kyoto University; Genome Integrity Unit, The Children’s Medical Research Institute, Westmead, NSW Australia; Department of Microbiology and Molecular Genetics, University of California, Davis; Graduate School of Biostudies and Radiation Biology Center, Kyoto University

**Keywords:** Chromatin modification, H2A.Z, H3K9me2, Trophoblast

## Abstract

During the transition from pluripotency to a lineage-committed state, chromatin undergoes large-scale changes in structure to effect the required changes to the transcriptional program. This involves covalent modification of histone tails, replacement of histone variants, and alteration in the subnuclear position of genes, including associations with the nuclear periphery. Here, using high-resolution microscopy and quantitative image analysis, we surveyed a panel of histone variants and covalent modifications for changes in nuclear periphery association during differentiation of human embryonic stem cells to a trophoblast-like lineage. This differentiation process is rapid and homogeneous, facilitating the use of a relatively fine timecourse (12h, 24h, and 48h post-initiation) to enable detection of transient changes. With this scheme, we detected two modifications with significant changes in enrichment at the nuclear periphery: acetylation of histone variant H2A.Z, and dimethylation of histone H3 at lysine 9. We show that these chromatin marks increase specifically at the nuclear periphery in a sequential, complementary manner, with a H2A.Z acetylation preceding H3K9 dimethylation. The increase of H3K9 dimethylation occurred coincidentally with but independently of accumulation of Lamin A, since Lamin A^-/-^ hES cells showed no changes in the localization pattern of H3K9 dimethylation. Inhibition of histone deacetylases led to persistent and increased H2A.Z acetylation at the periphery, and failure to differentiate. Our results show that a concerted dynamic change in the nature of peripheral chromatin is required for differentiation into the trophoblast state.

## Introduction

Pluripotent cell differentiation is accompanied by extensive changes in the expression of pluripotency and lineagespecific genes. Gene expression changes are controlled in turn by several mechanisms which include differential activity of transcription factors, covalent modifications of DNA or chromatin, and movement of gene loci into silent or active nuclear compartments (1). While our understanding of the transcriptional networks required for the stem cell state and commitment to a variety of lineages is well developed, (2) our knowledge regarding the role of chromatin changes and nuclear subcompartments in differentiation is not as comprehensive. Some clues have emerged from work on mouse embryonic stem cells (mESCs), where upon differentiation, genes which are required in the newly established cell lineages shift from being late replicating to earlier replicating (3, 4). This shift in replication timing also correlates with dissociation from the nuclear periphery during differentiation, since DNA at the periphery tends to be replicated later than active, euchromatic regions (1, 5). The nuclear periphery is in general a gene-repressive compartment (6); however, no universal causal relationship between gene position and transcriptional state has been defined. Movement of genes to and from the nuclear periphery is influenced by the covalent modification of histone proteins (7); especially, association of heterochromatin with the nuclear periphery is strongly correlated with dimethylation of histone H3 at lysine 9 (8–11). H3K9me2-marked chromatin can undergo dynamic relocalization to and from the nuclear periphery in a lineage-specific manner during differentiation (12). The nuclear lamina, a meshwork at the nuclear periphery, is composed of Lamin A, its splice variant Lamin C, Lamin B, and their interacting proteins. Of these, Lamin A/C and the Lamin B Receptor (LBR), have been strongly implicated in anchoring heterochromatin to the nuclear periphery in multiple cell types (13). While Lamin A and its splice variant Lamin C make up the bulk of the nuclear lamina in differentiated cells, these are only expressed at low levels in pluripotent cells; consequently, the lamina in pluripotent cells is composed largely of only Lamin B. Whether the low level of Lamin A present in undifferentiated human ES cells plays a role in transcriptional control is not clear.

Several reports have shown that the differentiation of stem cells is accompanied by significant changes in covalent DNA and histone protein modification which influence the activity of genes (14, 15). H3K9 di- or trimethylation is consistently associated with transcriptional repression, and targets lineage-associated genes during differentiation (16). Large H3K9me2 domains are present in both undifferentiated and differentiated cells (17), but these undergo dynamic changes at the local level during differentiation. H2A.Z is a histone variant whose many roles are modulated by post-translational modifications, and is exchanged for canonical H2A by the Tip60 complex. While H2A.Z itself has been found in both repressive and activating contexts (18), its acetylated form has exclusively been found correlated with gene activation (19–21). Global and predictable changes in chromatin modifications between different nuclear compartments in differentiated and undifferentiated stem cells have also been reported (22). Despite these studies, we still know little about the kinetics of establishing chromatin environments during lineage commitment. Especially, the relationship between chromatin modifications and different nuclear compartments in cells at intermediate states between pluripotency and terminal differentiation is not understood well. Further, while information from ensembles of cells is able to reveal information about chromatin changes at high genomic resolution, it fails to show changes in 3D nuclear architecture of individual cells.

Trophoblast cells are the first cells to terminally differentiate *in vivo*, and eventually make up the extraembryonic placental tissue. Trophoblast-like (TBL) cells can be effectively and homogeneously differentiated from primed hESCs through treatment with BMP4 and simultaneous inhibition of FGF and Activin / Nodal signaling (23, 24). Compared to other commonly employed differentiation pathways to generate neural progenitor or mesoderm lineages, TBL differentiation generates cells that are immediately terminally committed, finish differentiation in less than 1 week, and exhibit characteristic chromatin states, including significant hypomethylation and distinct chromatin organization in a highly homogeneous manner (25–27). The fast, unidirectional, homogeneous nature of TBL cell differentiation provides a tool to study cellular events that happen immediately upon differentiation.

In this study we have analysed individual human embryonic stem cells (hESCs) to assay the localization of modified histones with respect to the nuclear periphery during early phases of pluripotency loss and into the beginning stages of terminal lineage commitment to trophoblast-like (TBL) cells. We visualized histone modifications with immunofluorescence using high resolution microscopy, and quantitatively analyzed the three-dimensional nuclear distribution of histone modifications relative to the nuclear periphery. To this end, we developed an automated program which recognizes and crops individual nuclei in three dimensions, and executes shell analysis to quantitatively examine distance-dependent distribution patterns of histone modifications relative to the nuclear periphery in hundreds of cells. Our shell analysis program is applicable to many different types of cells and signals whose distribution relative to nuclear periphery are to be determined. We surveyed dynamic changes of modified chromatin (H3K4me3, H3K9me2, H3K9me3, H3K27ac, H3K27me3, H3K36me3) and one histone variant (H2A.Z) and its acetylated form (H2A.Zac) in hESCs from both primed and “naïve” pluripotent states (28, 29) in addition to hESCs which are differentiating to the trophoblast lineage. We show that two histone modifications, the dimethylation of histone 3 at lysine 9 (H3K9me2) and N-terminal acetylation of the histone variant H2A.Z (H2A.Zac) show changes in their distribution at the nuclear periphery as hESCs transition from naïve to primed and differentiated states. Particularly, we have captured transient association of H2A.Zac to the nuclear periphery within 12h upon loss of pluripotency, and found that this coincides with the transient loss of H3K9me2 from the nuclear periphery during the differentiation process. Our analysis uncovered that transient and dynamic nuclear periphery association of specific histone modifications may occur during intermediate stages of differentiation. Such transient changes could be missed by only analyzing cells after the completion of differentiation. Further, we show that HDAC activity is a key controlling factor in the establishment of chromatin environments at the nuclear periphery during differentiation.

## Results

### Characterization of fast, homogeneous differentiation from hES cells to the TBL lineage

We first characterized the extent to which our TBL induction scheme could produce homogeneous, rapid pluripotency loss and differentiation from hES to TBL cells. We subjected primed hESCs to a treatment inducing TBL differentiation, as previously described (23, 24) (**Fig. 1A**, see Methods) and observed changes in gross cellular morphology, as well as the development of characteristic chromatin morphology revealed by DAPI staining (**Fig. 1B**). This DAPI morphology is qualitatively similar to that seen in early human placental sections (27). Immunofluorescence indicated that TBL induction caused a drop in NANOG and OCT4 (POU4F1) levels within 24 hours (**Fig. 1 C**), and single-molecule RNA FISH confirmed that *NANOG* mRNA levels were downregulated at this time (**Fig. 1 D**). The mRNA for *CDX2*, a gene essential for placental development (30), was detectable 48 hours post induction and was specifically detected in a subset of cells at 96 hours that displayed DNA morphology qualitatively similar to *in vivo* human cytotrophoblast (27) (**Fig. 1 E**). Immunofluorescence also demonstrated that the trophoblast specific marker Cytokeratin 7 was expressed in cells possessing the cytotrophoblast-like DNA morphology at 4 and 6 days post induction (**Fig. S1A**), as were other human trophoblast markers GCM1, SYNA and hCG (**Fig. S1B**). Lamin A, a nuclear envelope protein which is not abundantly expressed in pluripotent hESCs (31) and shows extremely sparse staining in undifferentiated hESCs, became clearly detectable at the nuclear periphery 48 hours after TBL induction (**Fig.1F**). Taken together, these results indicate our differentiation scheme is capable of rapid and homogeneous differentiation as measured by these cell fate markers.

**Fig. 1.**
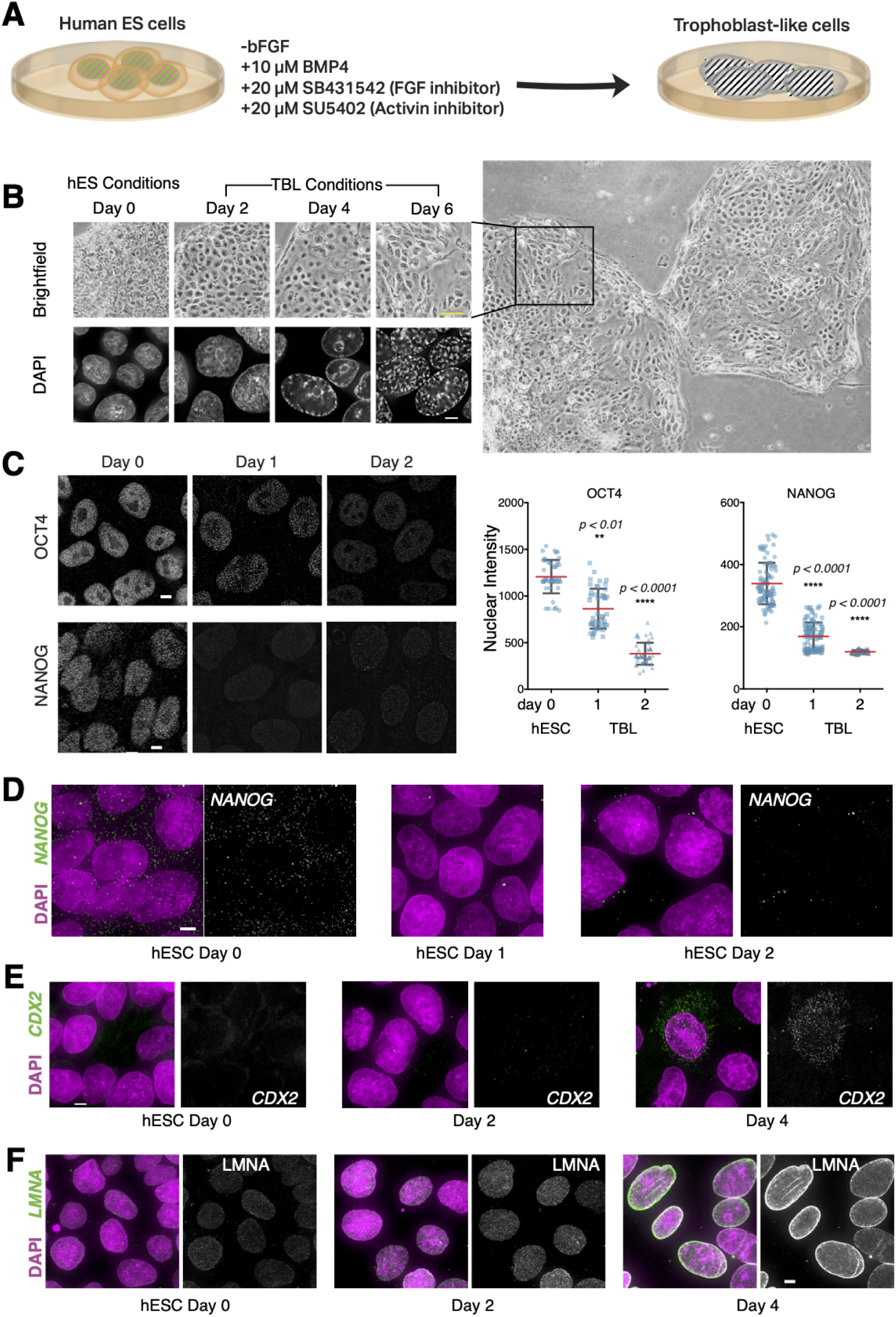
Changes in morphology and gene expression of human embryonic stem cells during differentiation to a trophoblast-like state. **A**, schematic of the differentiation procedure. **B**, Changes in colony morphology, nuclear staining using DAPI, and pluripotency markers Oct4 and NANOG using immunofluorescence, during differentiation. For the brightfield image of Day 6, the larger field of which it is a subset is shown adjacent. Scale bars: 30µm (top), 5µm (bottom). **C**, quantitation of intensity levels of Oct4 and NANOG from panel B; red lines show mean intensity values and bars extend to one standard deviation; significance levels for difference of means from day 0 are shown. **D**, single-molecule FISH against NANOG mRNA is shown in green; DAPI staining in magenta. **E**, single-molecule FISH against Cdx2 mRNA is shown in green; DAPI staining in magenta. **F**, immunofluorescence against Lamin A is shown in green; DAPI staining in magenta. Scale bars, 5µm.

### H2A.Zac and H3K9me2 undergo waves of relocation at the nuclear periphery as hESCs lose pluripotency and differentiate

We reasoned that if both subnuclear position and covalent histone modifications were involved in modulating gene expression, then subnuclear changes in histone modification should be cytologically visible across a differentiation time course. Using immunofluorescence, we examined the intensity and localization pattern of several histone modification marks associated with either active or inactive chromatin (H3K4me3, H3K9me2, H3K9me3, H3K27ac, H3K27me3, H3K36me3) as well as one histone variant (H2A.Z) and its acetylated form (H2A.Zac), in hESC and at 24 hours post TBL induction (**Fig. 2**). The subnuclear distribution of H3K4me3, H3K36me3, H3K27me3 and H3K9me3 did not noticeably change following 24 hours of differentiation (**Fig. 2 C, D, F** and **H**). The subnuclear distribution of pan-H2A.Z and H3K27ac within the nucleus also remained similar throughout differentiation, although nuclear intensity qualitatively decreased in differentiating cells (**Fig. 2 A, E**).

**Fig. 2.**
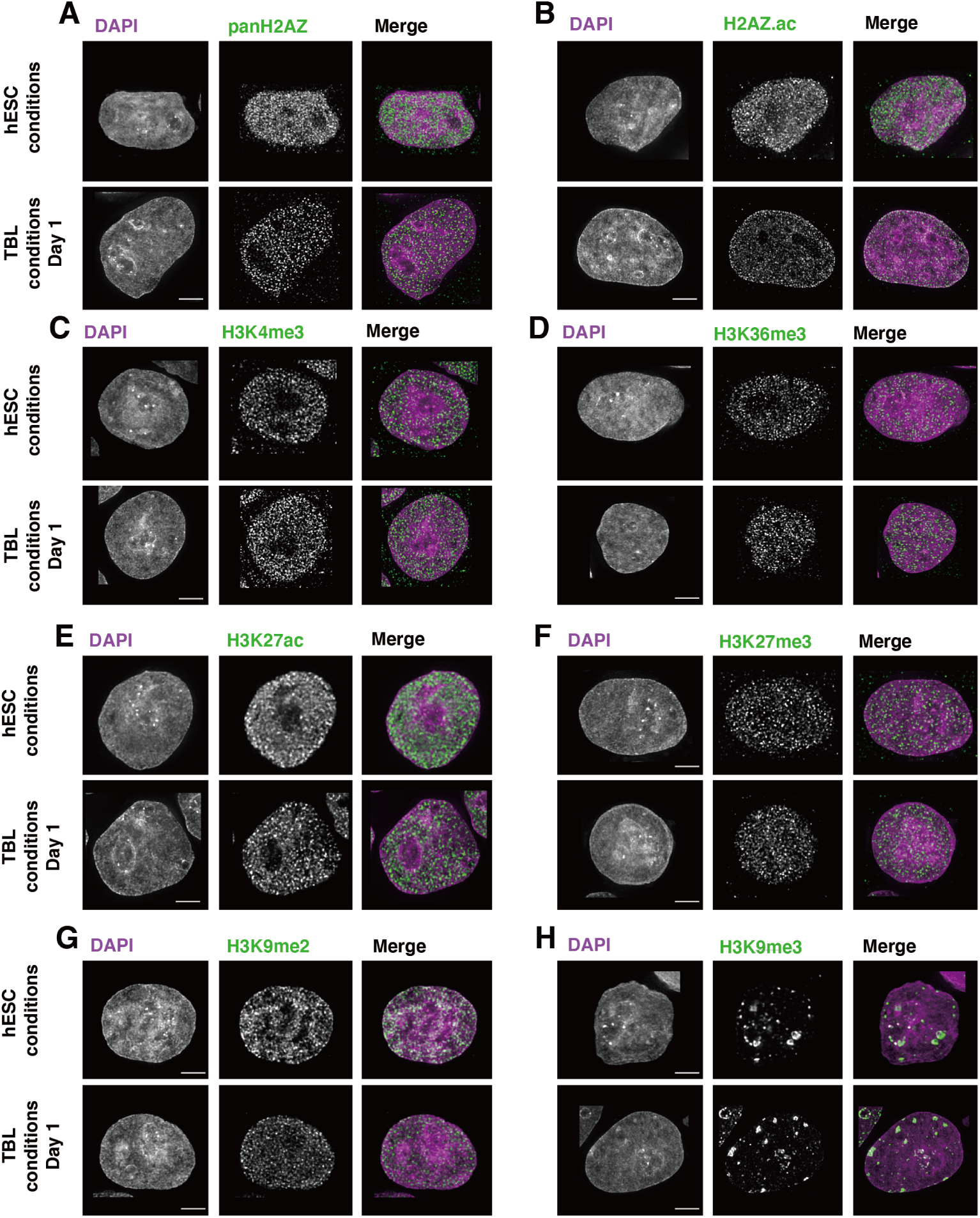
Chromatin markers display dynamic changes during differentiation. Cells in either hES media (top) or 24h after changing to TBL differentiation media (bottom) were stained with antibodies against the indicated chromatin markers. DAPI staining is shown in grayscale and magenta in the merged column; chromatin markers shown in grayscale and green in the merged column. Scale bars, 5µm.

In contrast, two of the tested histone modifications, H2A.Zac and H3K9me2, showed marked changes in nuclear distribution as cells lost pluripotency. Under hESC conditions, H2AZac was distributed evenly throughout the nucleus. After 24 hours of differentiation, H2AZac signal appeared enriched at the nuclear periphery and was reduced in the nucleoplasm (**Fig 2B**). Interestingly, H3K9me2 signals appeared to change in a manner opposite to that of H2AZac: undifferentiated hESCs displayed enriched H3K9me2 at the nuclear periphery, which was lost 24 hours after differentiation (**Fig 2G**).

To fully characterise H2A.Zac and H3K9me2 dynamics across all hESC stages, we took samples at 12, 24, 48 and 96 hours after the commencement of differentiation. To determine whether the pattern of histone modifications in non-differentiated hESC is characteristic of the primed state and the naïve state (a distinct, developmentally-earlier state of pluripotency in which cells are less prone to differentiate), we compared conventionally-cultured hESC (referred to hereafter as primed hESC) to naïve hESC. To quantitatively assess the apparent enrichment of signals at the nuclear periphery by immunofluorescence, we implemented a shell averaging program operating on three-dimensional images (**Fig. 3A**). The source images for the program are three-dimensional fields of ∼15µm depth containing from 2-10 cell nuclei. The DAPI channel is used to automatically segment each nucleus into its own new image, and is also used to define the nuclear periphery. The mid-section of the image is automatically detected, and fluorescence intensity is then averaged in each of a series of twenty concentric shells reaching from 0.5µm outside the nuclear periphery to ∼1.5µm inside, in five Z sections surrounding the midsection. The shells intentionally begin outside the true nuclear periphery to create a detectable peak in the nuclear signal. For each experiment, a montage of all segmented nuclei is automatically generated, which is used to visually exclude instances of failed segmentation (e.g., partial nuclei; or several nuclei being falsely detected as a single nucleus).

**Fig. 3.**
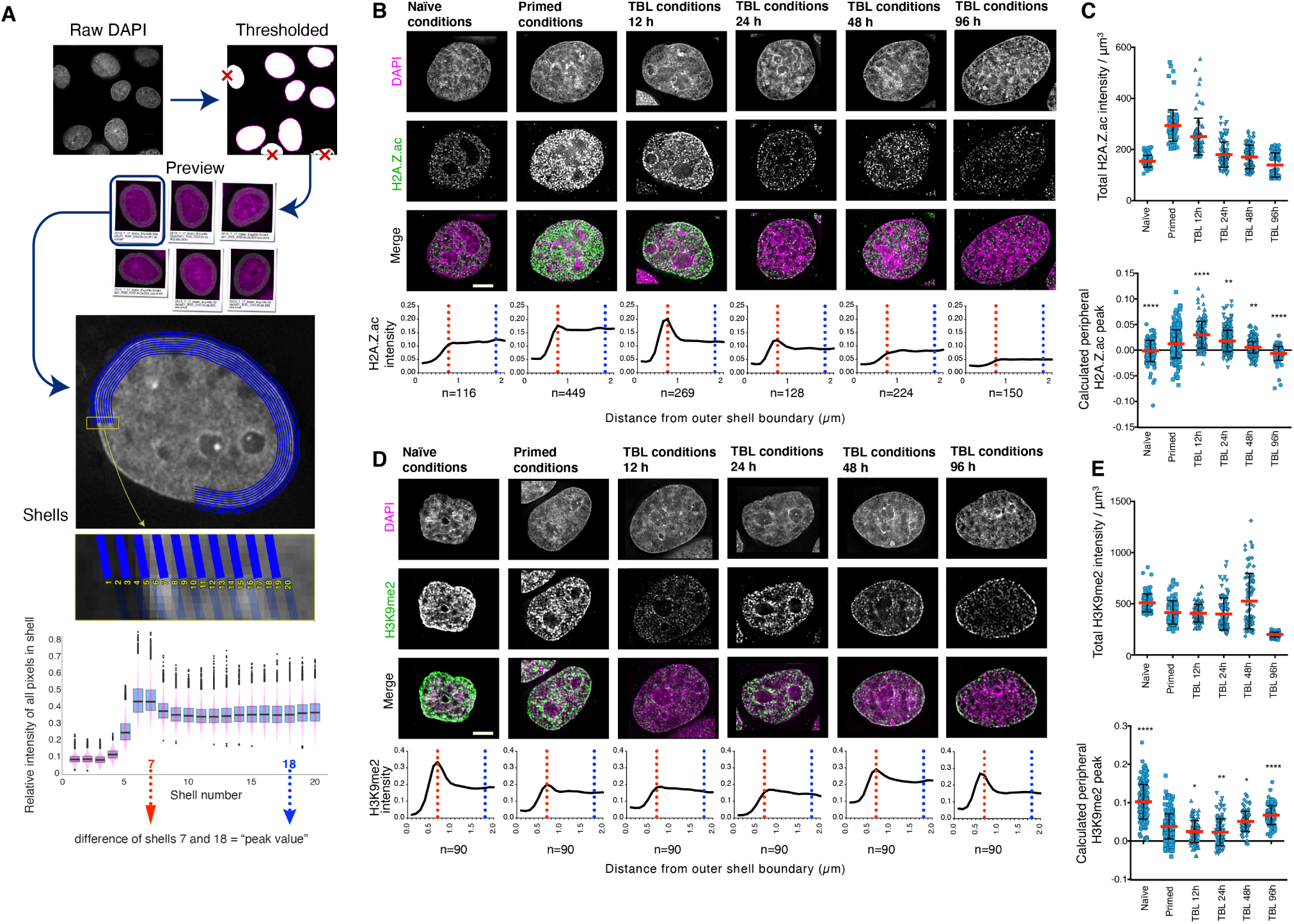
H2A.Z acetylation and H3K9me2 are alternately enriched at the nuclear periphery during differentiation. **A**, illustration of the shell-averaging procedure. The DAPI image is used to define a set of concentric shells spanning the nuclear periphery. The intensity of chromatin markers within each shell is averaged for the number of cells indicated underneath each graph; the difference between shells 7 and 18 is used to calculate the peak value. **B**, immunofluorescence examples of H2A.Zac under different differentiation conditions (top) and plots of shell-averaged intensity; cells and graphs shown are those nearest the average peak value for their class. **C**, plots of total intensity (top) and peak values (bottom) of H2A.Zac for all cells: n=(116, 449, 269, 128, 224, 150) from left to right. **D**, immunofluorescence examples of H3K9me2 and corresponding lots, under the same conditions shown in B. **E**, plots of total intensity (top) and peak values (bottom) of H3K9me2 for all cells: n=90 cells for all conditions. Scale bars, 5µm

For each experiment, cells were fixed at different time points and kept at 4°; immunostaining of all samples was then conducted at the same time to minimize technical variability. Images were acquired with the same exposure time in each wavelength to facilitate direct comparisons, and at least 90 cells total were analyzed from two or three biological replicates in all cases. The reliability of this method was confirmed by a consistent 200nm average peak distance measured between the peak of Lamin A/C immunostaining and the peak of DAPI staining in differentiated cells (**Fig. S2**). Using this method, we found that H2A.Zac was initially low and diffuse in naïve hESC, but in primed hESC, H2A.Zac was found more strongly throughout the nucleus and was occasionally enriched at the nuclear periphery (**Fig. 3B**). At 12 hours post TBL induction, most hESCs showed peripheral enrichment of H2AZac while internal levels declined. After 24 hours of differentiation cells still exhibited H2A.Zac enrichment at the nuclear periphery, but both peripheral and nucleoplasmic levels appeared reduced compared to cells at 12 hours. At 2 days post differentiation, most nuclei no longer displayed peripherally-enriched H2A.Zac, and by 96 hours H2A.Zac signal became very low throughout the whole nucleus. Quantitative analysis showed that total nuclear H2A.Zac signal was low in naïve hESC, high in primed hESCs and generally declined with differentiation (**Fig. 3C)**. Peripheral shell analysis across the population demonstrated that there were some cells with peripheral H2A.Zac enrichment at all timepoints assayed, but at 12 and 24 hours post differentiation the majority of the population exhibited peripheral H2A.Zac.

To test whether peripheral enrichment was specific to the acetylated version of H2A.Z, we also characterised panH2A.Z dynamics. In both naïve and primed hESC pan-H2A.Z was seen as small nucleoplasmic foci that were present throughout the nucleus, though more intense in primed cells (**Fig. S3**). After 24 hours of TBL induction, pan-H2A.Z levels remained high in the nucleoplasm, but further differentiation was accompanied by a decrease back to naïve levels. While pan-H2A.Z detected by immunofluorescence also changed dynamically, it was not enriched at the nuclear periphery, indicating that the periphery is specifically enriched for acetylation of H2A.Z.

We next assayed H3K9me2 dynamics across the 4-day differentiation time course. We found that H3K9me2 appeared at the nuclear periphery in a pattern strikingly opposite to H2A.Zac. H3K9me2 was significantly peripheral in naïve cells, reaching peak intensity within 0.7 µm from the thresholded nuclear periphery (**Fig. 3D**), the same shell as the peak of DAPI intensity due to peripheral heterochromatin. Peripheral enrichment was reduced in primed hESC relative to naïve hESC nuclei, and by 12 hours following the initiation of differentiation, both peripheral and internal H3K9me2 signal were significantly reduced (**Fig. 3D**). Nucleoplasmic H3K9me2 signal returned at 24 hours post differentiation and from 48 hours, cells exhibited robust H3K9me2 signal at the nuclear periphery (**Fig. 3D**). Quantitative analysis showed that the total H3K9me2 signal appeared relatively consistent across differentiation (**Fig. 3E**). Population wide peripheral shell analysis revealed that primed hESC and cells at 12 and 24 hours of differentiation displayed reduced H3K9me2 signal at the nuclear periphery relative to populations of both naïve cells and cells that had been differentiating for 48 hours or more (**Fig. 3F**).

We next investigated whether any correlation existed between the levels of H2A.Zac and H3K9me2 at the nuclear periphery, and how these changes related to pluripotency loss. We performed simultaneous immunostaining of H2A.Zac and H3K9me2 with Lamin A. We located cells that still possessed some peripheral enrichment for H2A.Zac at 24 hours post differentiation, and compared their peripheral profiles to neighboring cells in the same image that had undergone H2A.Zac reduction. We found a positive correlation between H2A.Zac reduction, H3K9me2 accumulation, and differentiation status as judged by the intensity of the peripheral Lamin A border (**Fig. 4**).

**Fig. 4.**
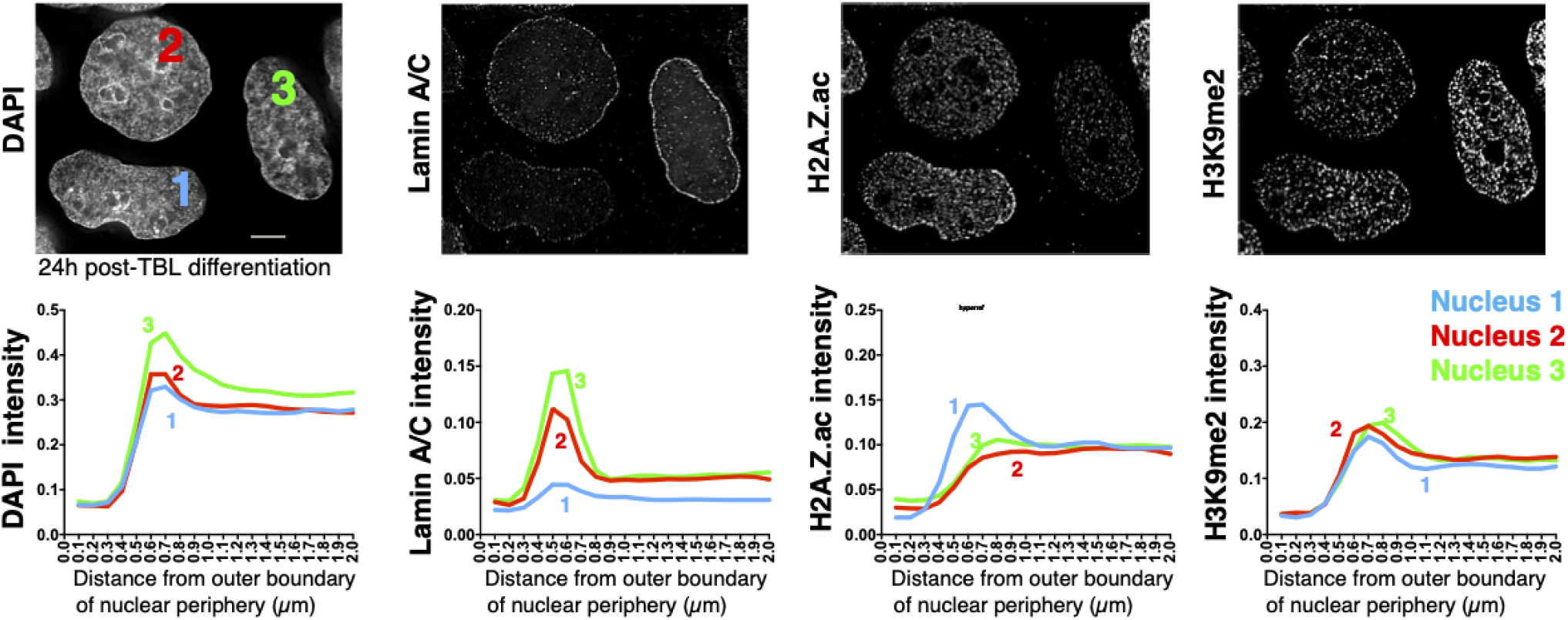
Comparison of individual cells at the same differentiation timepoint shows alternation of H2A.Zac and H3K9me2 peripheral intensity. Cells shown stained with DAPI and immunofluorescence as indicated; shell-averaged intensity graphs are shown for each numbered cell. Scale bar, 5µm.

### Lamin A is not necessary for peripheral chromatin reorganisation during hESC differentiation

Given that gradual expression and localization of Lamin A at 48 hours after TBL differentiation coincides with re-appearance of H3K9me2 at the periphery, we hypothesized that Lamin A might be a factor influencing the dynamic changes of H3K9me2 localization to the nuclear periphery. To test this, we used CRISPR to generate a hESC line with a homozygously disrupted *Lmna* gene (**Fig. 5A**). *Lmna*^*-/-*^ hES cells proliferated normally and did not manifest any morphological differences compared to control cells under normal culturing conditions (**Fig. S4)**. Immunofluorescence microscopy showed a complete loss of Lamin A staining in knockout cells, in contrast to the typical nuclear rim staining visible in unedited cells (**Fig. 5B**). Despite the loss of Lamin A, no qualitative differences in H3K9me2 localization or intensity could be seen between homozygous knockout and unedited cells upon TBL induction (**Fig. 5C, D**). *Lmna*^*-/-*^ hES cells were able to differentiate into other lineages, including beating cardiomyocytes and neural progenitor cells (NPC) by directed differentiation induction (**Supplemental Movie 1** and data not shown). We therefore conclude that although Lamin A localization in hESC is highly correlated with H3K9me2 at the periphery, it is not required for its localization.

**Fig. 5.**
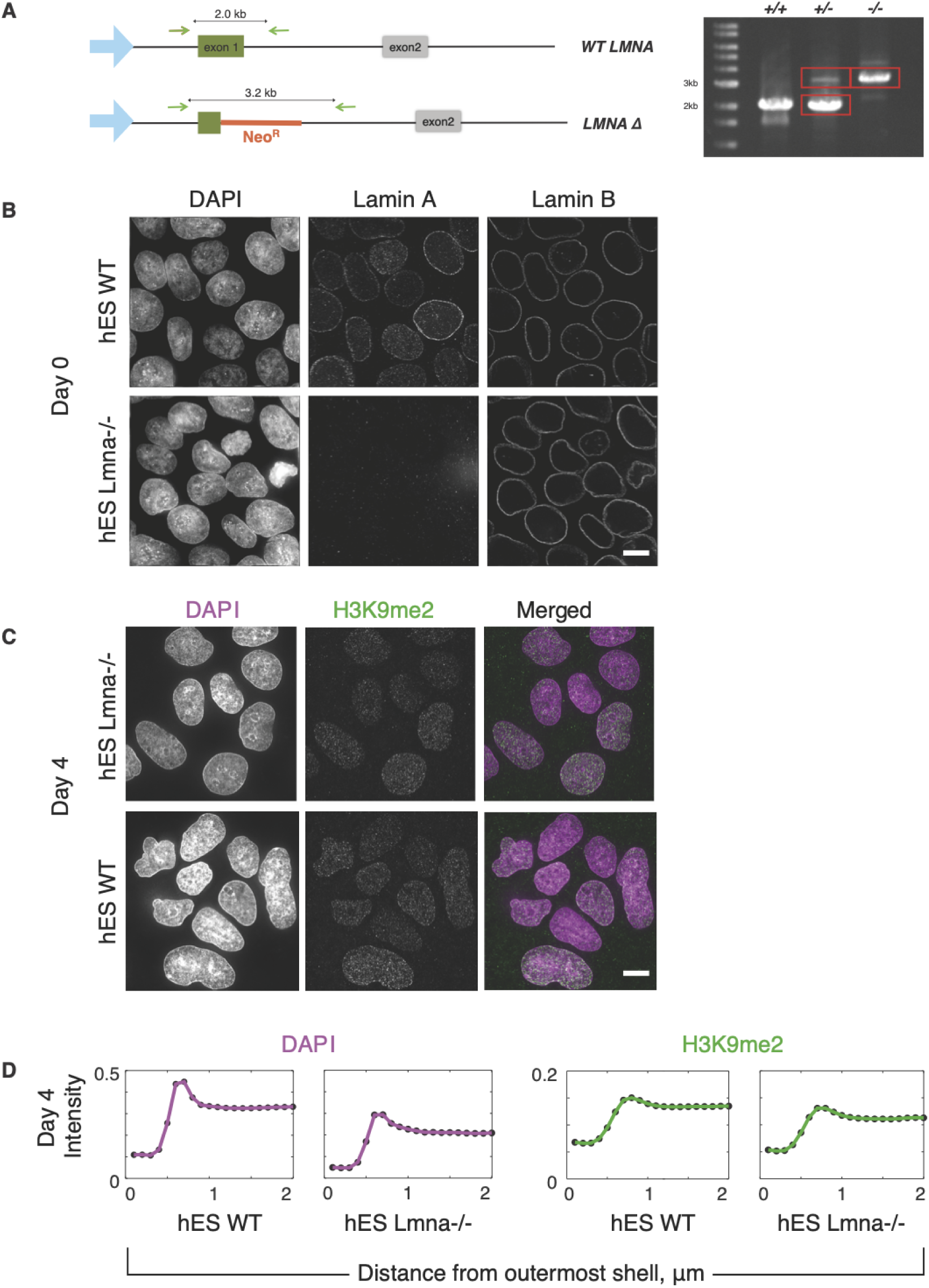
Lamin A does not play a role in H3K9me2 dynamics during TBL differentiation. **A**, Illustration of gene editing to remove the LmnA gene. CRISPR was used to replace the first exon of LmnA with a neomycin resistance cassette, disrupting the gene; hetero- and homozygous deletions were detected by PCR as shown (right). **B**, WT hES cells (top) and Lmna^-/-^ hES cells (bottom) stained for DAPI, Lamin A, and Lamin B. **C**, DAPI and H3K9me2 staining in WT hES cells (top) and Lmna^-/-^ hES cells (bottom) 4 days post-TBL differentiation. **D**, Quantitation of DAPI and H3K9me2 peripheral shell staining in WT hES cells and Lmna^-/-^ hES cells as indicated. Scale bars, 5µm.

### The localization of HAT and HDAC proteins is dynamic as hESCs differentiate

We next wondered how H2A.Zac levels were changed by acetylation (i.e. histone acetyl-transferase (HAT) activity) versus deacetylation (i.e. histone deacetylase (HDAC) activity). We reasoned that if histone modifications were enriched at the nuclear periphery, then the enzymes responsible for them may also be cytologically detectable in the same location. Previous work suggests that Tip60 is the primary HAT catalyzing H2A.Z acetylation (20, 32, 33), and while the identity of HDACs responsible for deacetylation of H2A.Zac is not known, it is likely to be a member of the “Class I” HDACs, HDAC1, HDAC2 or HDAC3 (34). HDAC1 and HDAC2 are known to be required for self-renewal and differentiation in ES and TS cells; however, the role of HDAC1 seems to be more critical (35–37).

We found that in primed hESC, Tip60 existed in 3 primary patterns, as central nucleoplasmic clusters surrounding DAPI enriched areas; as diffuse nucleoplasmic foci with no clear enrichment pattern; or, as nucleoplasmic foci with a tendency for enrichment at the nuclear periphery (**Fig. S5C**). Each specific Tip60 pattern could be found in cells at 12h and 24h post-TBL induction as well; however, the proportion of nuclei possessing different localisations changed, with primed hESC showing greater tendency toward peripheral localization than naïve or differentiating cells. In contrast, HDAC proteins displayed mostly homogeneous nuclear localization by immunofluorescence. We found that both HDAC1 and HDAC2 were localized to the nucleus in hESC. HDAC2 was stained intensely in hESC nuclei (**Fig. 6A**), while HDAC1 presented as small foci throughout the nucleoplasm (**Fig. S5A**). HDAC3 was not significantly enriched in the nucleus of pluripotent hESCs but did become nuclear 48 hours after cells were subjected to TBL induction (**Fig. S5B**). HDAC1 levels appeared to drop as hESCs began to differentiate (**Fig. S5A**) while HDAC2 levels were retained throughout differentiation, with some increased signal at 12 and 24 h post TBL induction (**Fig. 6B**). Of the HDACs we surveyed, therefore, the expression pattern of HDAC2 most closely tracks the loss of H2A.Zac at the periphery.

**Fig. 6.**
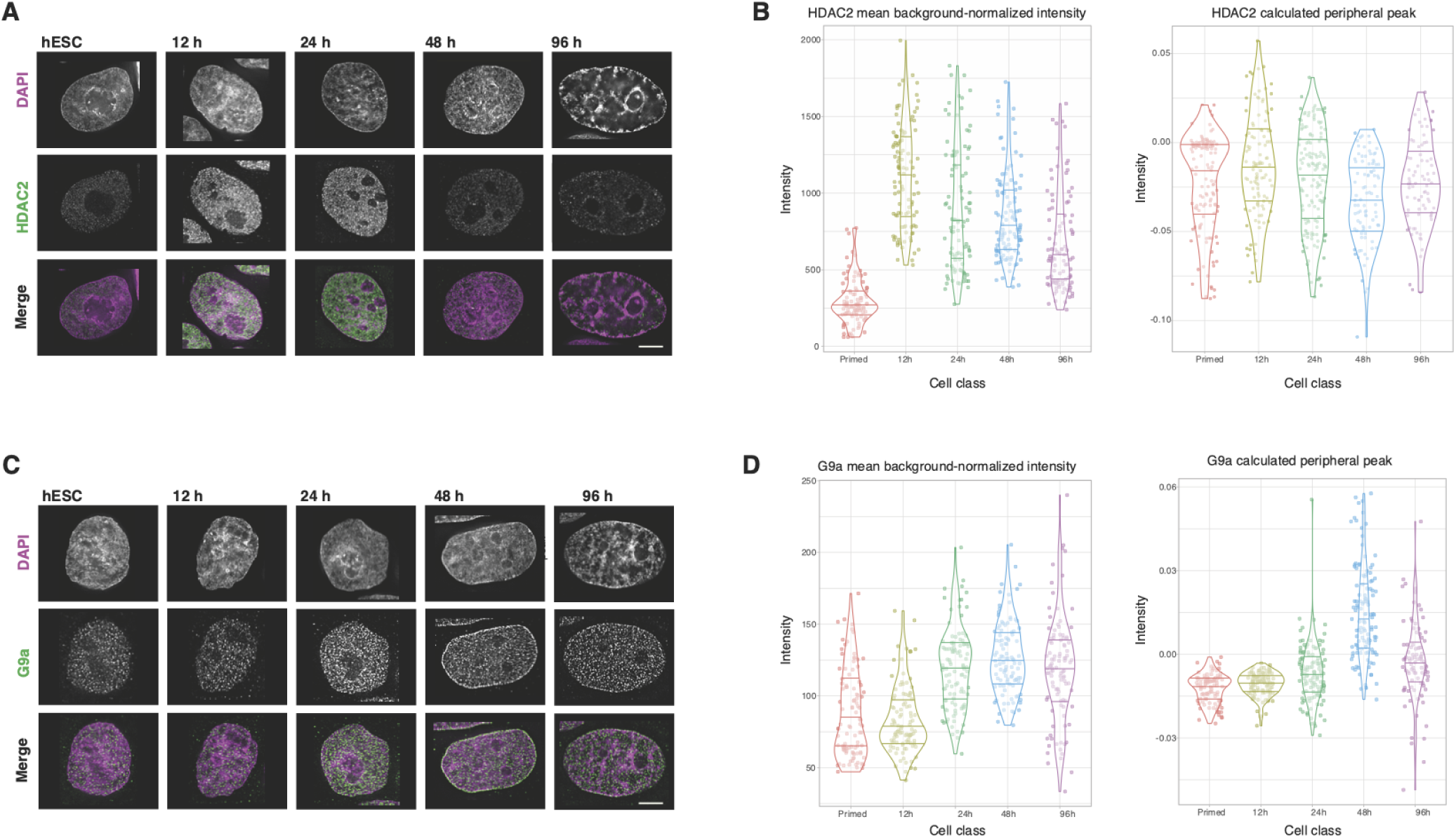
HDAC2 and G9a display dynamic localization during TBL differentiation. **A**, Representative images of HDAC2 immunostaining during a TBL differentiation time course. **B**, quantitation of mean background-corrected intensity of HDAC2 for all nuclei at each timepoint (left) as well as the peak value calculated by subtracting the mean of shell 18 from shell 7 (right). All data points are shown as dots; distribution density shown by violin plot. n=(104, 102, 101, 112, 98) cells measured from left to right. **C**, representative images of G9a immunostaining during TBL differentiation. **D**, mean background-corrected intensity of G9a for all nuclei at each timepoint (left) as well as the corresponding peak values (right). For both HDAC2 and G9a, the image chosen for each timepoint is closest to the mean value for total intensity. n=(100,103,108,105,104) cells measured from left to right. Scale bars, 5µm.

The modifier specifically responsible for catalyzing H3K9 dimethylation is the histone methyl-transferase (HMTase) G9a (38). In primed hESC, G9a was detected by immunostaining as small nuclear foci throughout the cell nucleus (**Fig. 6C**). G9a location tracked that of H3K9me2 from 24 hours post differentiation, with a significant increase in G9a intensity throughout the nucleoplasm and a measurable increase in G9a signal at the nuclear periphery in hESC at 24 and 48 hours post differentiation (**Fig. 6D**). Together, our data suggests that HAT, HDAC and HMTase complexes responsible for changes in H2A.Z acetylation status and H3K9 methylation status also exhibit dynamic changes in their localization during pluripotency loss and early phases of differentiation.

### Acetylation of H2A.Z and dimethylation of H3K9 at the nuclear periphery is driven by histone modifiers and is essential for the progression of differentiation

We next asked whether pharmacologically inhibiting histone modification enzymes would affect the localization of histone marks, and subsequently pluripotency loss and/or differentiation. We performed a series of experiments using the HDAC inhibitor TSA, a global inhibitor of Class I HDACs (39) expected to inhibit HDAC1, HDAC2 and HDAC3. Cells were first subjected to TBL differentiation, and then exposed to 12.5nM TSA for either 12h or 36h before sampling; control cells without TSA were sampled in parallel (**Fig. 7A**). Inhibition with TSA is reversible, and does not alter HDAC protein levels nor association, only deacetylation kinetics (40). This allowed us to indirectly interpret gains in histone acetylation in different nuclear compartments as locations where Class I HDAC complexes would usually be active (**Fig. 7B**). To determine if HDAC activity drives H2A.Zac removal from the nuclear periphery during differentiation, hES cells were induced to differentiate and treated with TSA at 12 hours, when peripheral H2A.Zac is elevated yet global levels have begun to decline. Under these conditions, H2A.Zac remained enriched at the nuclear periphery and H3K9me2 signal at the periphery did not increase even after 48 hours of growth under differentiation conditions (**Fig. 7C**,**D)**.Lamin A at the nuclear periphery was also less intense in treated cells, suggesting that TSA-treated hESC did not differentiate (**Fig. 7E)**. These results suggest that HDAC activity, which appears to drive H2A.Zac removal from the nuclear periphery, is required for the normal progression of differentiation.

**Fig. 7.**
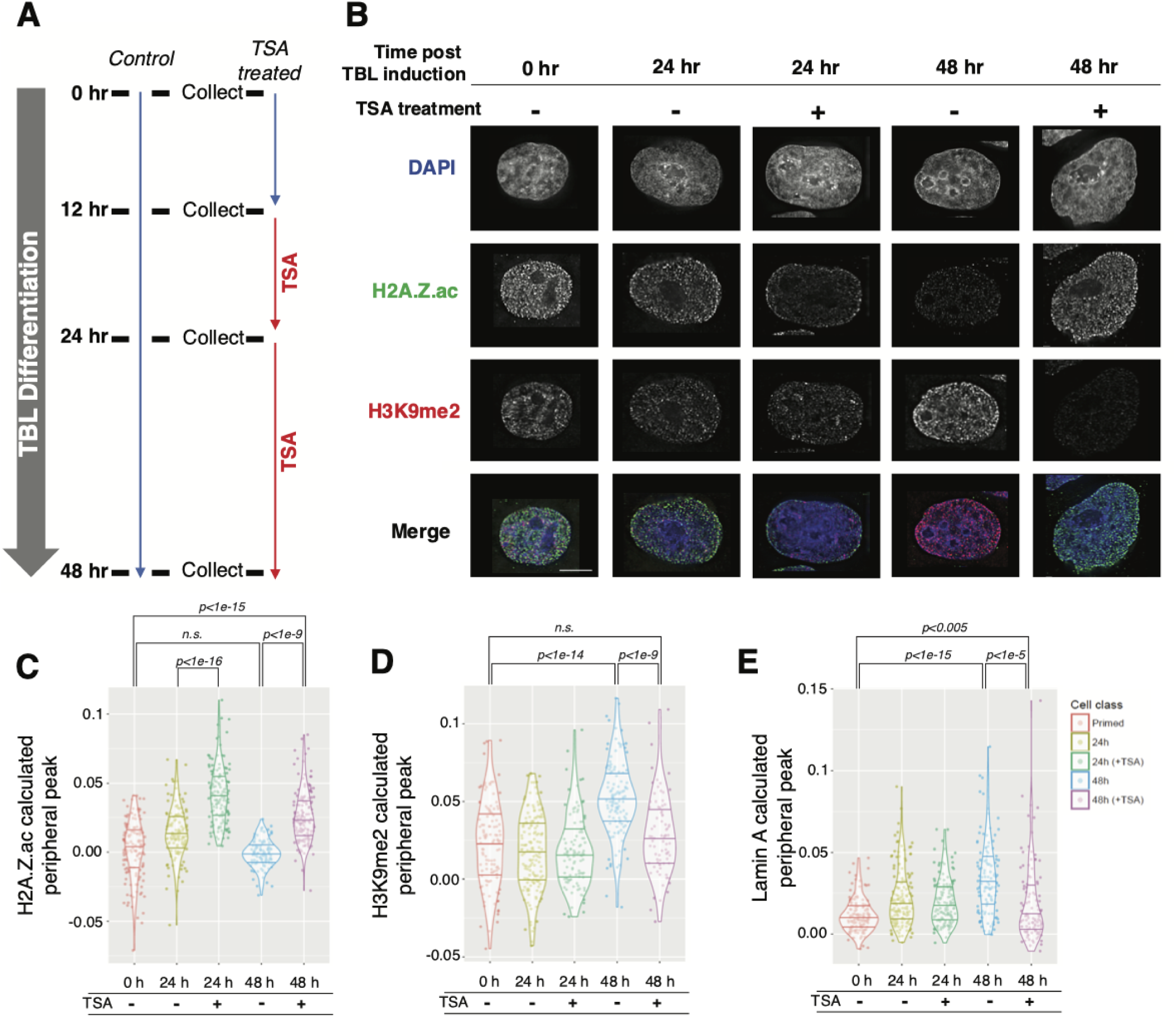
TSA treatment prevents differentiation-associated changes in H2A.Z acetylation. **A**, schematic of treatment and collection times. TBL differentiation begins at time t0. **B**, representative images of nuclei immunostained for H2A.Zac and H3K9me2. **C**, peak values of H2A.Zac calculated for individual cells; n=142, 128, 141, 101, 149 from left to right. **D**, peak values of H3K9me2 calculated for individual cells: n=116, 153, 102, 121, 81 from left to right. **E**, peak values of Lamin A calculated for individual cells: n=134, 153, 129, 122, 111 from left to right. Scale bars, 5µm. Significance values assessed with two-tailed t test.

## Conclusions

By quantitatively measuring the intranuclear distribution of histone modifications and variants during TBL differentiation, we find that H2A.Zac and H3K9me2 are the most significantly altered. H2A.Zac transiently elevates at the nuclear periphery of hESCs just prior to loss of pluripotency, but is then lost; whereas from day 2 of differentiation, H3K9me2 becomes peripherally elevated instead. We have shown that ongoing HDAC activity is responsible for removing H2A.Zac from the nuclear periphery at the onset of differentiation, and that HDAC activity is essential for differentiation to proceed. From analysis of the transcriptional activation and repression of the androgen receptor system (21, 33), Giaimo (18) have proposed that H2A.Z, and especially ubiquitinated H2A.Z, can be repressive, but H2A.Z acetylation leads to its deubiquitination and consequent activation of genes. Our finding that the nuclear periphery is specifically and transiently enriched for H2A.Z acetylation in the early stages of pluripotency loss of human ES cells suggests that upon differentiation, many genes normally found at the periphery, a generally repressive compartment, may rapidly transit to an active state through the action of H2A.Z-specific acetyltransferases. While the current study does not identify the genes that may be so regulated, it predicts that many genes whose activation is regulated by H2A.Z acetylation during TBL differentiation are likely to be found in hES cell Lamin-associated domain (LAD) compartments.

Other studies have assayed the subnuclear distribution of histone modification marks, including H3K4me2 and me3, H3K9me1, me2 and me3, H3K9ac, H3K27me2 and me3 and H3K79me1, in pluripotent and differentiated hESCs induced to differentiate by non-directed means. These studies reported that the only histone modifications which exhibited changes in radial distribution were H3K9me3, H3K9ac and H3K79me1, which were described as being reduced at the nuclear periphery (22). However our results showed that H3K9ac and H3K9me3 showed no significant location differences as hESCs lose pluripotency. These discrepancies could arise from differences in differentiation protocols and fate (our TBL protocol versus non-directed differentiation) as well as the image analysis methods used to calculate peripheral enrichment. In our method, we use an algorithm to define the periphery of the nucleus based on thresholded DAPI staining, and then assign shells by distance working inwards, as compared to the method employed by (22), wherein radial shells were established working outwards from the nucleus center. It is likely, however, that the particular histone marks enriched in the periphery vary, as commitment to different lineages involves different signals and transcriptional programs.

Treating primed hESC with the HDAC inhibitor TSA caused persistent and increased intensity of H2A.Zac throughout the nucleoplasm and at the nuclear periphery. This suggests that there is constant, active de- and re-acetylation in the pluripotent ES cell, including at the periphery. In addition to persistent H2A.Zac at the periphery, H3K9me2 appearance at the periphery was correspondingly delayed in TSA-treated cells, suggesting that the two histone modifications are regulated by linked mechanisms that trigger upon pluripotency loss.

In conclusion, we have shown evidence that the redistribution of two chromatin modifications (H2AZac and H3K9me2) at the nuclear periphery is a hallmark of pluripotency loss and/or differentiation. We have shown that Lamin A is not necessary for the alternation of H2A.Zac and H3K9me2 at the periphery during TBL differentiation. We have also shown through cytological analysis that H2A.Zac in general negatively correlates with H3K9me2. We suggest that in the naïve hESC, H3K9me2 exists at the periphery to reinforce the peripheral location of silenced genes. However, when the hESC cell enters a primed state, Tip60 (or other H2A.Z acetylase) activity is higher at the periphery, enabling H2A.Z to be rapidly acetylated at the periphery following HDAC 1 and/or HDAC2 reduction upon pluripotency loss. This could be to prevent a premature accumulation of H3K9me2 at the nuclear periphery, thereby ensuring the proper restructuring of chromatin upon lineage commitment and the refinement of key regulating elements such as LADs. Future work using single-cell ChIP-Seq analysis of histone marks could determine the identity of the genes regulated by the peripheral changes we observe and give important clues into compartment-specific gene regulation during differentiation.

## Methods

### Cell Culture

Primary mouse embryonic fibroblast (MEF) cells were grown under standard conditions (5% CO2 humidified environment at 37°C) in DMEM media supplemented with 10% fetal bovine serum. Human ES cells (H1 line) were maintained in ReproCell Primate media supplemented with FGF2 StemBeads (8µL / mL, Stem Cultures) on Mitomycin-C inactivated MEFs and mechanically passaged every 5-7 days. Following methods described in (28), hESCs were induced to a naïve state by growing cells in “naïve Human Stem Cell Media” (NHSM) which was ReproCell Primate media containing LIF (20 ng / mL), TGFβ (1ng / mL), FGF2 (8 ng / mL), PD0325901 (1µM), CHIR99021 (3µM), SP600125 (10µM) and SB203580 (10µM) (28). NHSM media was added to hESCs 3 days post passage, and cells were allowed to grow for 2 days before a subsequent passage. Cells were then grown for a further 5 days in NHSM and used as “naïve” hESCs after the next passage. To differentiate hESCs into trophoblast like (TBL) cells, hESCs were passaged onto MEF-free surfaces coated with ES grade matrigel (Fisher Scientific, Cat# 354277) and grown in ReproCell FF2 media and supplemented with FGF2 StemBeads. TBL differentiation was induced by removing hESC media and growing cells with recombinant BMP4 (10 ng / mL), SB431542 (20µM) and SU5402 (20µM). For TSA treatments, TSA was added to media to a final concentration of 12.5 nM, 12 hours after the start of differentiation induction. Cardiomyocyte differentiation of WT and *Lmna*^*-/-*^ hES cells was performed using the inducing agent KY02111 according to protocols in (41).

**Table 1.**
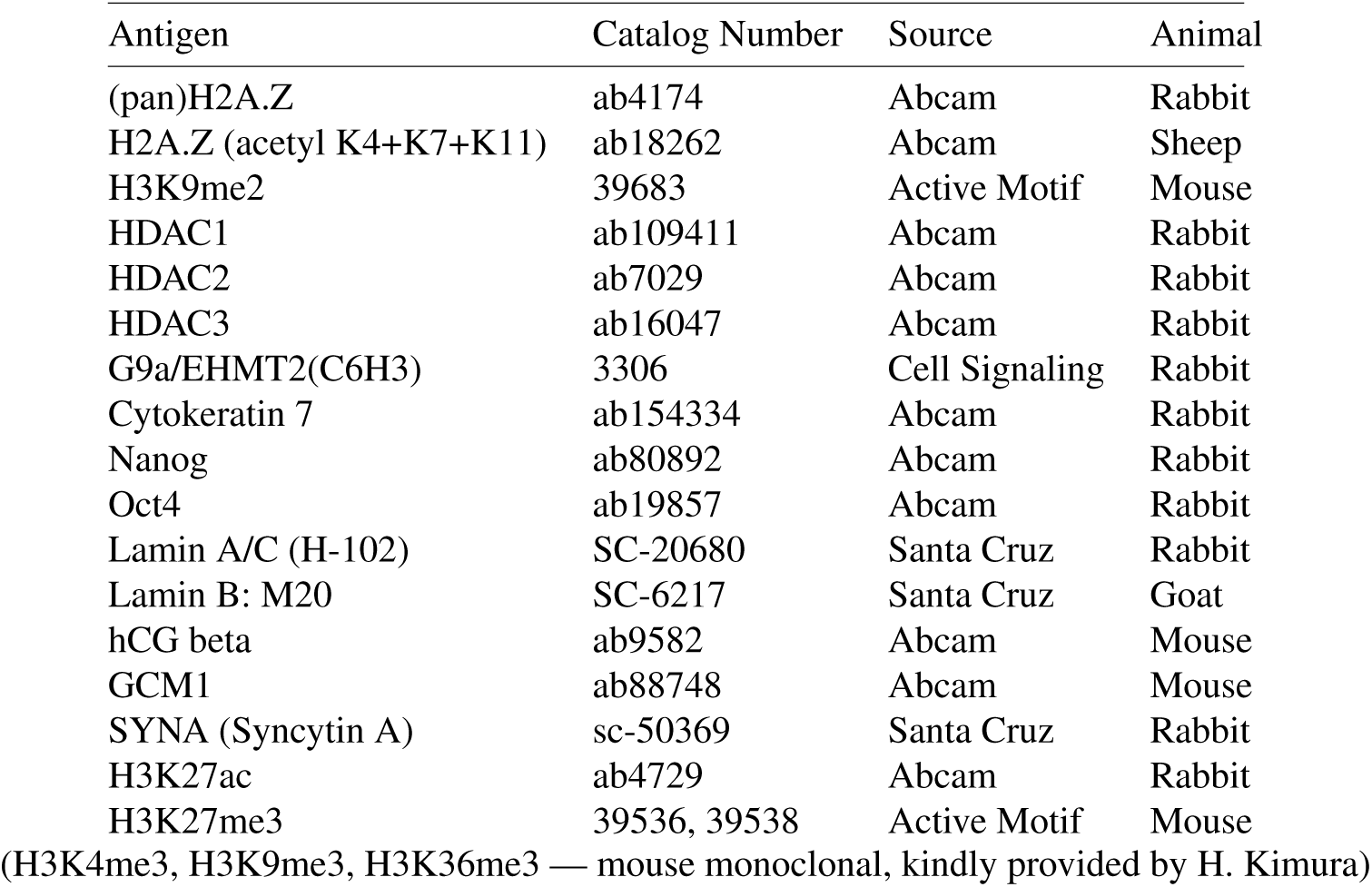
Antibodies used

| Antigen | Catalog Number | Source | Animal |
| --- | --- | --- | --- |
| (pan)H2A.Z | ab4174 | Abcam | Rabbit |
| H2A.Z (acetyl K4+K7+K11) | ab18262 | Abcam | Sheep |
| H3K9me2 | 39683 | Active Motif | Mouse |
| HDAC1 | ab109411 | Abcam | Rabbit |
| HDAC2 | ab7029 | Abcam | Rabbit |
| HDAC3 | ab16047 | Abcam | Rabbit |
| G9a/EHMT2(C6H3) | 3306 | Cell Signaling | Rabbit |
| Cytokeratin 7 | ab154334 | Abcam | Rabbit |
| Nanog | ab80892 | Abcam | Rabbit |
| Oct4 | ab19857 | Abcam | Rabbit |
| Lamin A/C (H-102) | SC-20680 | Santa Cruz | Rabbit |
| Lamin B: M20 | SC-6217 | Santa Cruz | Goat |
| hCG beta | ab9582 | Abcam | Mouse |
| GCM1 | ab88748 | Abcam | Mouse |
| SYNA (Syncytin A) | sc-50369 | Santa Cruz | Rabbit |
| H3K27ac | ab4729 | Abcam | Rabbit |
| H3K27me3 | 39536, 39538 | Active Motif | Mouse |

(H3K4me3, H3K9me3, H3K36me3 — mouse monoclonal, kindly provided by H. Kimura)

### Immunofluorescence

For experiments, cells were seeded onto glass coverslips or 35 mm glass bottom dishes (Matsunami#1S) coated with ES grade matrigel (Fisher Scientific, Cat# 354277) and fixed by incubation with 4% PFA for 10 minutes at 4 °C. Samples were washed with PBST and permeabilized with 0.5% Triton-X in PBS for 5 minutes at room temperature. Samples were washed again and incubated in blocking solution (2% BSA in PBST), then incubated with primary antibodies (**Table S1**) diluted in block solution in a humidified chamber overnight at 4°C. Samples were washed before incubation with secondary antibodies (**Table S1**) for 1 hour at room temperature, washed again and counterstained with DAPI for 5 minutes before being mounted onto microscope slides with 1% *n*-propyl gallate in glycerol. Samples were stored at −20°C prior to imaging.

### RNA FISH

Cells were grown glass coverslips and fixed as per the immunofluorescence procedure described above. FISH probes (Stellaris) directed against human Nanog and Cdx2 (see **Supplemental Data S1**) were reconstituted in nuclease-free TE buffer to a concentration of 25µM and stored at 20°C prior to use. Cells were permeabilized with cold 70% ethanol for 1 hour before hybridization, washed in wash buffer (10% formamide in 2X saline-sodium citrate (SSC) before hybridization was performed using probes diluted to 250 nM in hybridization buffer (10 mg/mL dextran sulfate, 10% formamide in 2X SSC) for 4 hours at 37°C. Cells were then washed again, nuclei were counterstained with DAPI in 2X SSC for 5 minutes before mounting onto microscope slides with 1% *n*-propyl gallate in glycerol. Samples were stored at −20°C prior to imaging.

### Fluorescence microscopy

Cells were imaged on a conventional wide-field DeltaVision deconvolution system (Applied Precision/GE Healthcare) using a 100x oil objective and immersion liquid of refractive index 1.513. Images were corrected for bleaching and lamp flicker, and processed with constrained iterative deconvolution using the softWoRx suite, using a point-spread function measured with 100nm green fluorescent latex beads at a Z step size of 0.2, to remove out-of-focus light. Axial aberration of different wavelengths was corrected through sub-pixel shifting in the Z direction based on peak axial positions of multicolor beads. Quantitative analysis of images was performed with the Fiji/ImageJ suite (42).

### Knockout of *LMNA* gene

hES cells (H1) were thawed at P29 and transfected with 3.3µM of each hCas9D10A (Addgene #74495), and a gRNA/donor plasmid (pUC_lmnA_Neo_exn1_donor_fixed containing gRNA sequence GCAGGAGCTCAATGATCGCT**TGG**; see **Supplemental data S2**) using Lipofectamine 2000 as per instructions. Cells were rescued and grown in the presence of ROCK inhibitor overnight, then placed in normal hES media on Matrigel for 4 days. Subsequently, G418 was added at 200µg/ml to select for Neo+ cells. After 2–3 days, cells were passaged to plates containing inactivated feeder MEF cells. After four days of colony growth, individual colonies were picked and grown in 24-well plates. Colonies were then sampled and screened by PCR for deletion. After establishment of the Lmna^-/-^ cell line, PCR and sequencing was carried out on a selection of sites in coding regions containing the nearest homology to the gRNA sequence, and no changes were found (data not shown). Primers used to screen for deletion were lmnA_nested_fwd [5’gggactgaagggggaagg3’] and lmnA_out_rev [5’cactctgcgtgtctggg3’], expected to give 2kb in WT cells but a 3.2kb band in deletion cells.

### Periphery distance analysis

For each nucleus, an absolute nuclear periphery was determined based on a smoothed, thresholded DAPI image. Calculating the Euclidean distance map inward from the threshold boundary and binning by 100nm (1.56 pixel increments) enabled the generation of 20 peripheral “shells” of 100nm each, beginning approximately 500nm outside the periphery and covering a total of 2 µm (see **Fig. 3A**). The average fluorescence intensity of Lamin A compares well with the peripheral distribution profile of DAPI staining, serving as an internal control for the shell analysis (**Fig. S2**). Code (GNU Octave and Perl) is provided at github.com/pmcarlton/shellanalysis.

### Statistical analysis

All statistical analysis was performed using R version 3.4.4 (43) or Prism (GraphPad, Inc.) software.

## ACKNOWLEDGEMENTS

We thank H. Kimura for kindly providing monoclonal antibodies, N. Nakatsuji for guidance and resources in maintaining human ES cells, I. Minami for assistance with cardiomyocyte differentiation, and K. Hasegawa and A. Sato for critical reading and helpful discussions. This project was carried out in part through funds provided by the Inamori foundation and the Sumitomo foundation.

---

**Supplemental Fig. 1.**
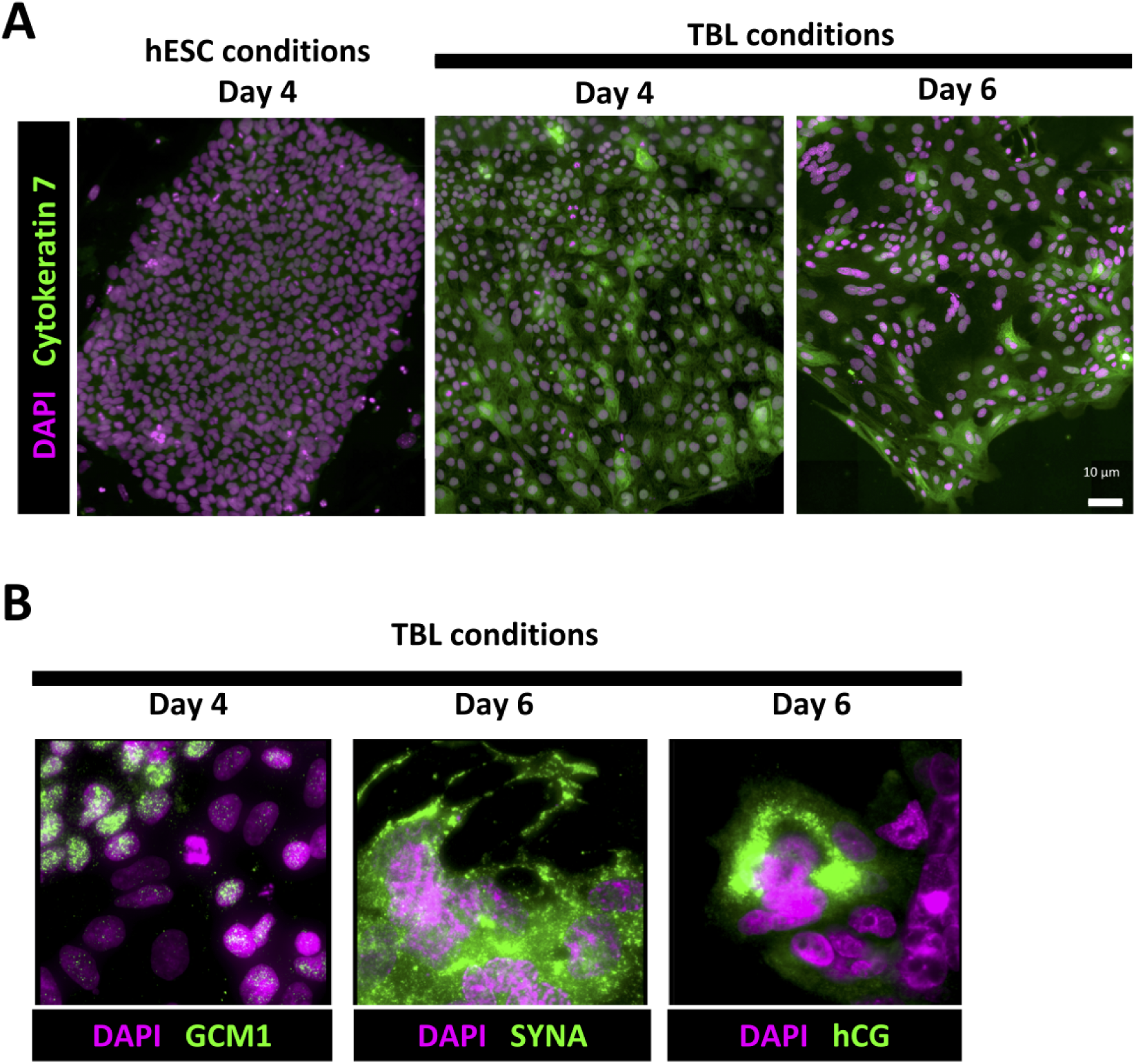
Immunofluorescence characterization of late-stage hES differentiation into a TBL fate. **A**, comparison of nondifferentiated (left) with differentiated day 4 (middle) and day 6 (right) cells, stained with DAPI (magenta) and Cytokeratin 7 (green). **B**, trophoblast biomarkers GCM1, Syncytin A, and hCG are immunostained in green.

**Supplemental Fig. 2.**
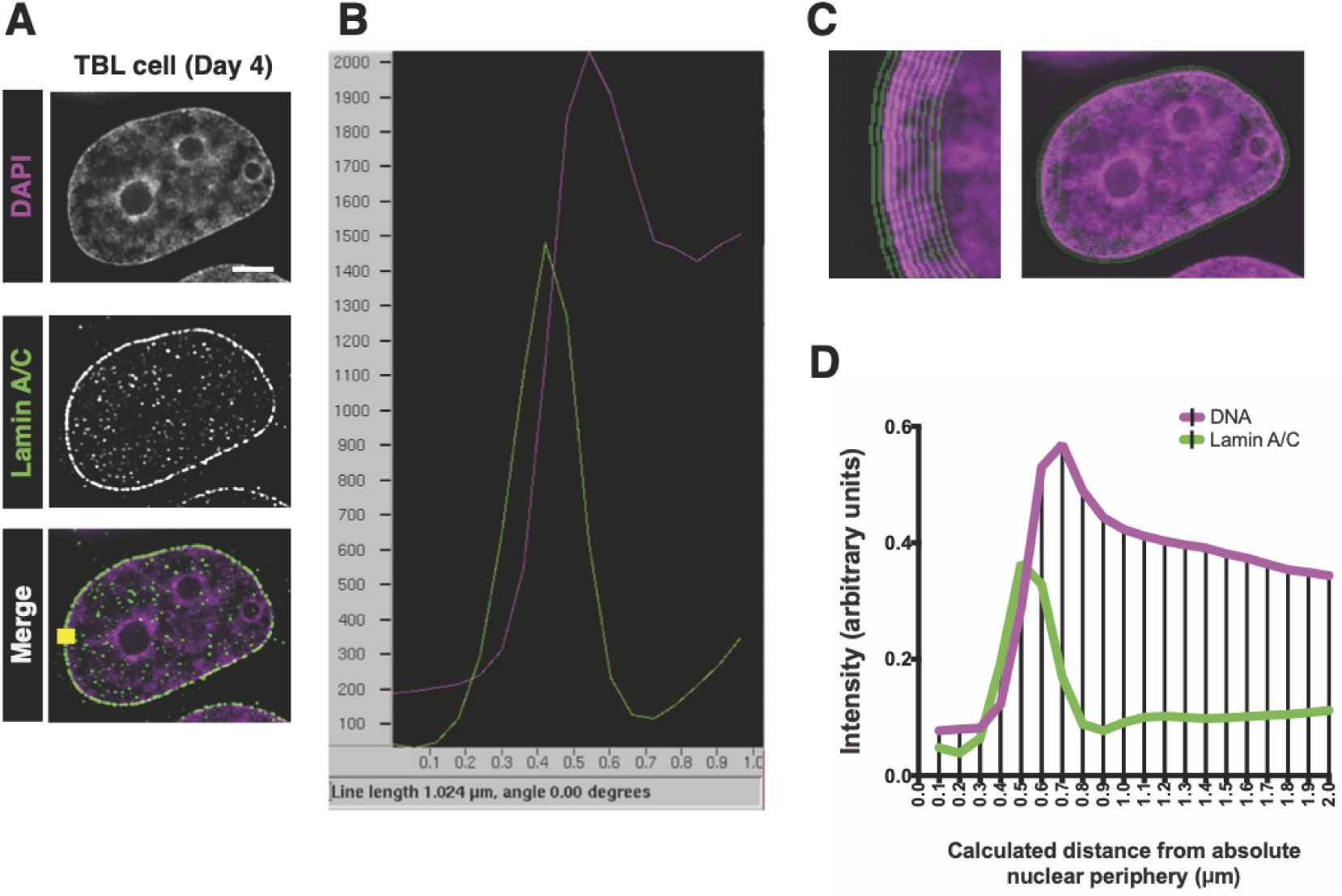
Illustration of shell averaging procedure and agreement with line profile. **A**, representative image of a hES-derived cell nucleus after 4 days of TBL differentiation treatment, stained with DAPI (top), Lamin A (middle); merged image at bottom. The bottom image shows a yellow line where the line profile is taken in panel B. **B**, a line profile showing intensity of both DAPI (magenta) and Lamin A (green) immunostaining. **C**, illustration of the 20 peripheral shells detected and used for intensity averaging. **D**, average intensities within each shell, calculated automatically, show agreement with the manually-drawn line profile.

**Supplemental Fig. 3.**
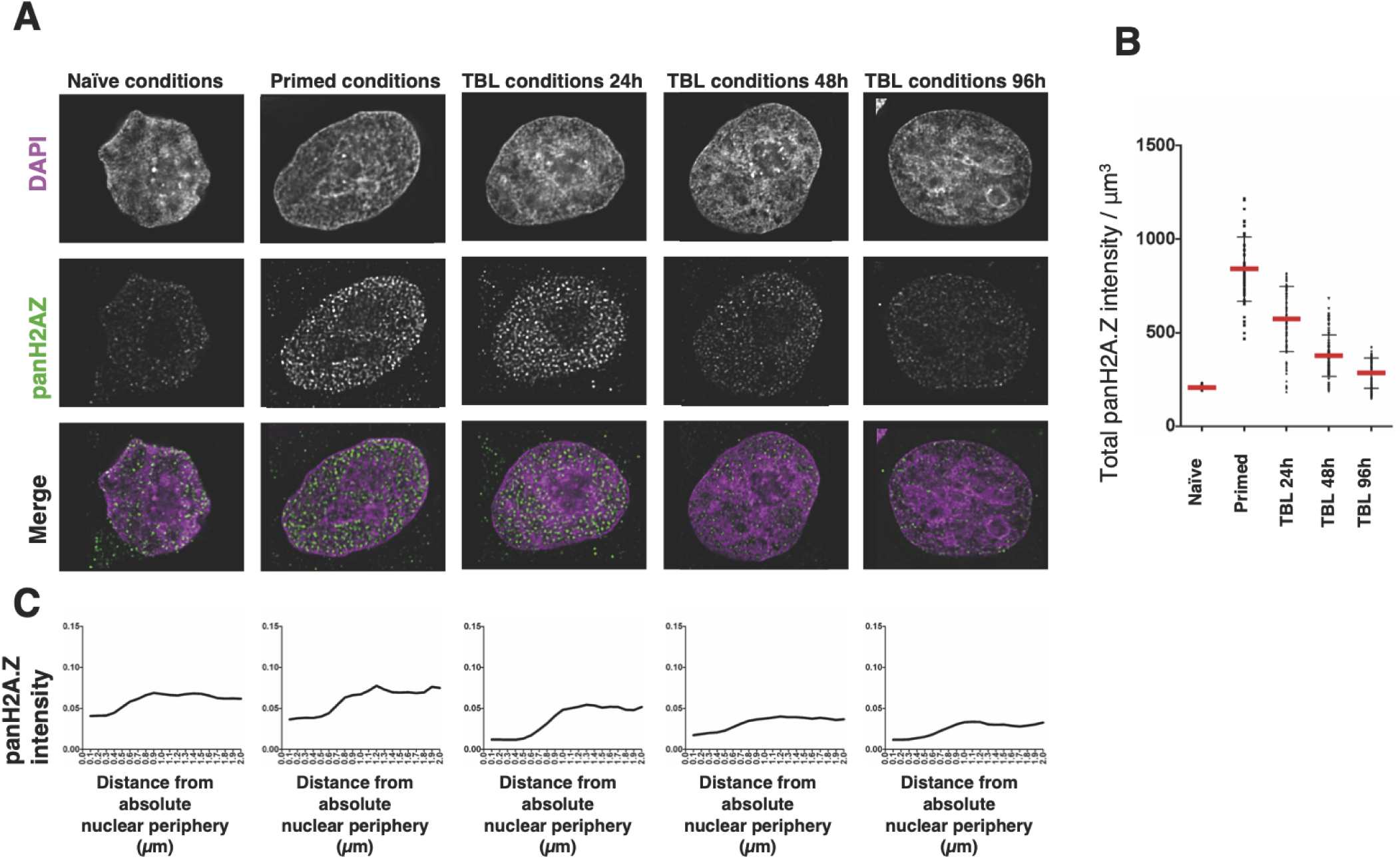
Immunofluorescence showing distribution of unmodified (pan-)H2A.Z. **A**, a timecourse of differentiation showing pan-H2AZ (middle row; green in merged image) and DAPI staining (top row, magenta in merged image). **B**, quantitation of H2A.Z intensity over the entire nucleus. **C**, peripheral shell analysis showing lack of enrichment at the nuclear periphery.

**Supplemental Fig. 4.**
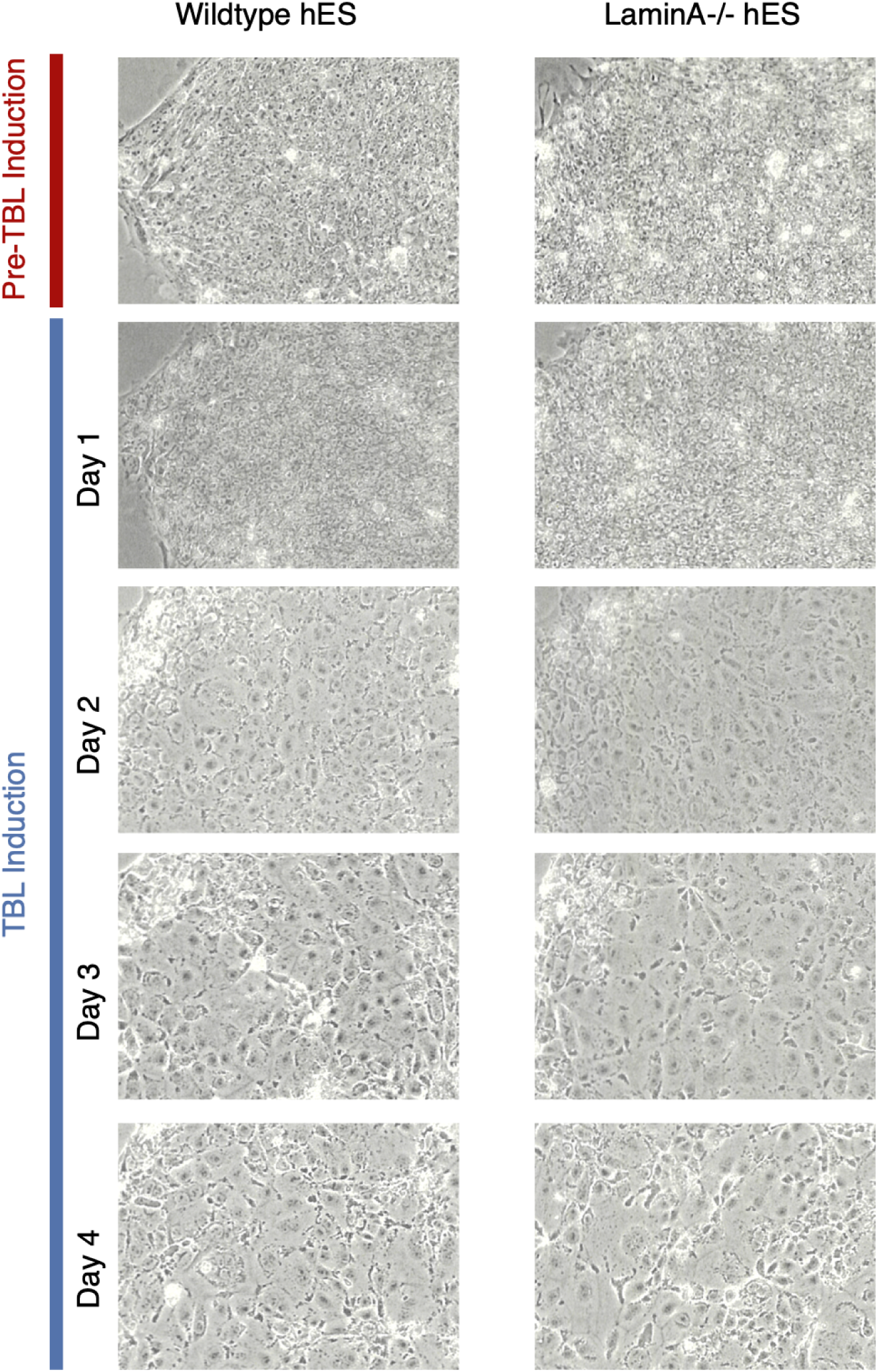
Bright-field fluorescence images of wild-type (left) and LmnA^-/-^ cells (right) during the TBL induction timecourse at indicated timepoints (all images shown at the same magnification).

**Supplemental Fig. 5.**
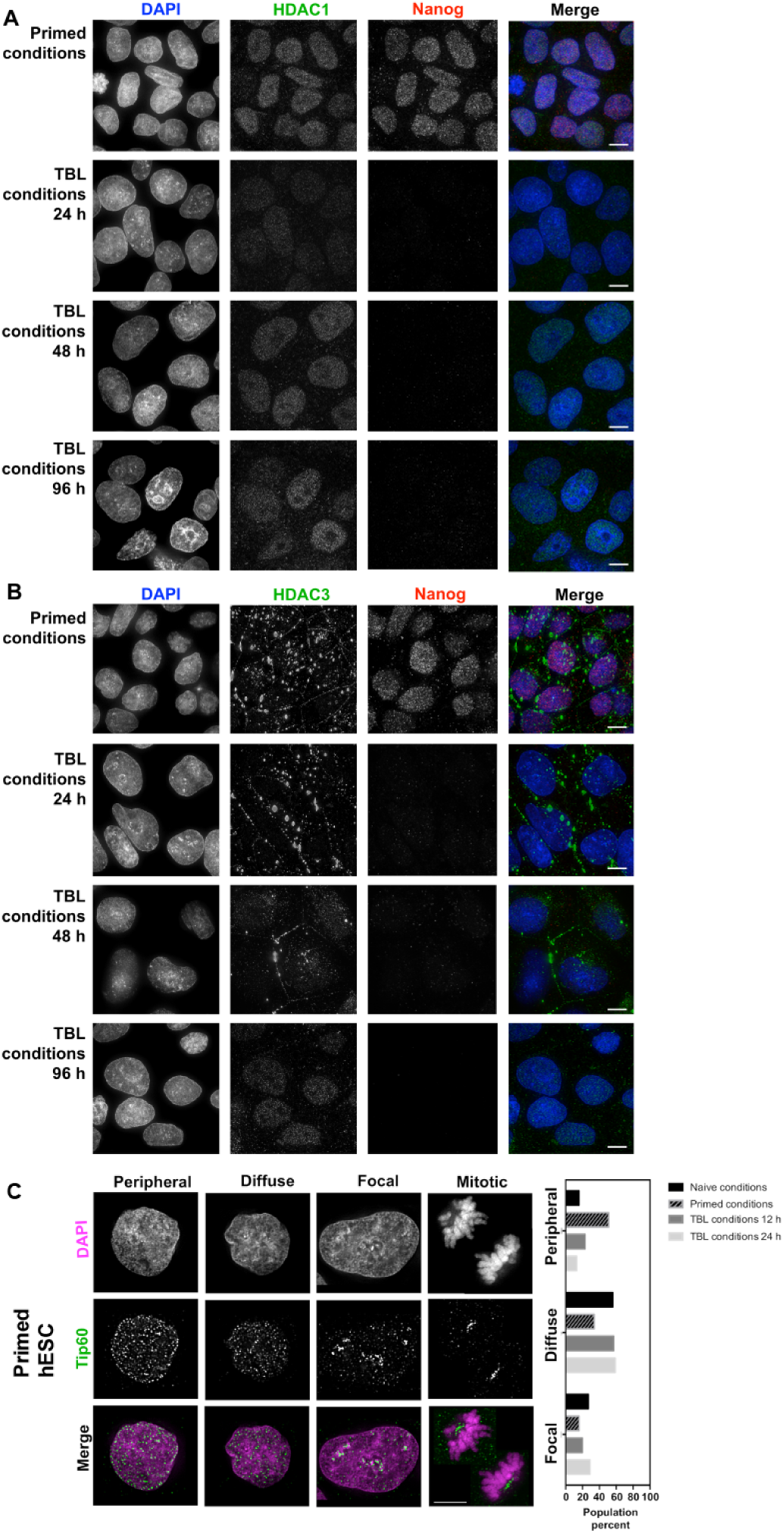
Immunofluorescence showing distribution of HDAC1 (**A**) and HDAC3 (**B**) over a differentiation timecourse. HDACs are labeled green in the merged image; Nanog immunofluorescence is labeled red in the merged image; DAPI counterstaining of DNA is shown in blue. Stages are indicated at left. Scale bars, 5µm. **C**, immunofluorescence of Tip60 staining in primed hESC. Representative images are shown for each staining pattern observed. *Right*, prevalence of each staining type as a fraction of population at the indicated times.

