## Supplemental Data S1 for "Sequential enrichment at the nuclear periphery of H2A.Zac and H3K9me2 accompanies pluripotency loss in human embryonic stem cells"

### Supplemental Data S1: Sequences (5'—3') for oligonucleotide RNA FISH

| Sequence | Sequence Name | Scale | Purification | Three Modificatic | MW Backbone |
|---|---|---|---|---|---|
| agtccagcagaacgttaaaa | nanog-pmc-mrna_1 | 30 nmol | Desalted | T(CAL Fluor Red | 7293.03 |
| aataacatgaggcaaccagc | nanog-pmc-mrna_2 | 30 nmol | Desalted | T(CAL Fluor Red | 7278.01 |
| ggataaagtgagttgcctgc | nanog-pmc-mrna_3 | 30 nmol | Desalted | T(CAL Fluor Red | 7347.04 |
| taggacctccagaaggaaaa | nanog-pmc-mrna_4 | 30 nmol | Desalted | T(CAL Fluor Red | 7318.04 |
| ggttaaggcagctttaagac | nanog-pmc-mrna_5 | 30 nmol | Desalted | T(CAL Fluor Red | 7331.04 |
| ggatccacactcatgttagt | nanog-pmc-mrna_6 | 30 nmol | Desalted | T(CAL Fluor Red | 7241.97 |
| aaggcaagctttggggacaa | nanog-pmc-mrna_7 | 30 nmol | Desalted | T(CAL Fluor Red | 7365.06 |
| ctttacagtcggatgcttca | nanog-pmc-mrna_8 | 30 nmol | Desalted | T(CAL Fluor Red | 7232.96 |
| aatcacaggcataggtgaag | nanog-pmc-mrna_9 | 30 nmol | Desalted | T(CAL Fluor Red | 7349.06 |
| ggatagttttcttcaggccc | nanog-pmc-mrna_10 | 30 nmol | Desalted | T(CAL Fluor Red | 7248.96 |
| agcagaagacattgcaagg | nanog-pmc-mrna_11 | 30 nmol | Desalted | T(CAL Fluor Red | 7349.06 |
| cagtctccgtgtgaggcatc | nanog-pmc-mrna_12 | 30 nmol | Desalted | T(CAL Fluor Red | 7258.95 |
| gctgtcctgaataagcagat | nanog-pmc-mrna_13 | 30 nmol | Desalted | T(CAL Fluor Red | 7291.01 |
| cgacactattctctgcagaa | nanog-pmc-mrna_14 | 30 nmol | Desalted | T(CAL Fluor Red | 7210.95 |
| gtttcttgactgggaccttg | nanog-pmc-mrna_15 | 30 nmol | Desalted | T(CAL Fluor Red | 7279.98 |
| ggaagagaacacagttctgg | nanog-pmc-mrna_16 | 30 nmol | Desalted | T(CAL Fluor Red | 7365.06 |
| cattgagtacacacagctgg | nanog-pmc-mrna_17 | 30 nmol | Desalted | T(CAL Fluor Red | 7275.99 |
| tgttggagagttcttgcatc | nanog-pmc-mrna_18 | 30 nmol | Desalted | T(CAL Fluor Red | 7304.01 |
| cctgtttgtagctgaggttc | nanog-pmc-mrna_19 | 30 nmol | Desalted | T(CAL Fluor Red | 7279.98 |
| cattctctggttctggaacc | nanog-pmc-mrna_20 | 30 nmol | Desalted | T(CAL Fluor Red | 7208.93 |
| tattcttcggccagttgttt | nanog-pmc-mrna_21 | 30 nmol | Desalted | T(CAL Fluor Red | 7229.96 |
| catccctggtggtaggaaga | nanog-pmc-mrna_22 | 30 nmol | Desalted | T(CAL Fluor Red | 7332.02 |
| gaaggttcccagtcgggttc | nanog-pmc-mrna_23 | 30 nmol | Desalted | T(CAL Fluor Red | 7298.98 |
| caggtctggttgctccacat | nanog-pmc-mrna_24 | 30 nmol | Desalted | T(CAL Fluor Red | 7233.94 |
| gttgctccaggttgaattgt | nanog-pmc-mrna_25 | 30 nmol | Desalted | T(CAL Fluor Red | 7304.01 |
| ctccaggactggatgttctg | nanog-pmc-mrna_26 | 30 nmol | Desalted | T(CAL Fluor Red | 7273.97 |
| caggtctgagtgttccagga | nanog-pmc-mrna_27 | 30 nmol | Desalted | T(CAL Fluor Red | 7323.01 |
| ctgattgttccaggattggg | nanog-pmc-mrna_28 | 30 nmol | Desalted | T(CAL Fluor Red | 7329.02 |
| tatagaagggactgttccag | nanog-pmc-mrna_29 | 30 nmol | Desalted | T(CAL Fluor Red | 7331.04 |
| ggactgcagagattcctctc | nanog-pmc-mrna_30 | 30 nmol | Desalted | T(CAL Fluor Red | 7242.95 |
| aaagcagcctccaagtcact | nanog-pmc-mrna_31 | 30 nmol | Desalted | T(CAL Fluor Red | 7204.94 |
| tgctgtattacattaaggcc | nanog-pmc-mrna_32 | 30 nmol | Desalted | T(CAL Fluor Red | 7256.99 |
| atccatggtttgtggagtac | nanog-pmc-mrna_33 | 30 nmol | Desalted | T(CAL Fluor Red | 7313.02 |
| catcttcacacgtcttcagg | nanog-pmc-mrna_34 | 30 nmol | Desalted | T(CAL Fluor Red | 7177.91 |
| ccagtgtccagactgaaatt | nanog-pmc-mrna_35 | 30 nmol | Desalted | T(CAL Fluor Red | 7250.98 |
| atcatggaaaccagaacacg | nanog-pmc-mrna_36 | 30 nmol | Desalted | T(CAL Fluor Red | 7278.01 |
| cctccatgagattgactgga | nanog-pmc-mrna_37 | 30 nmol | Desalted | T(CAL Fluor Red | 7266.98 |
| tgattaggctccaaccatac | nanog-pmc-mrna_38 | 30 nmol | Desalted | T(CAL Fluor Red | 7210.95 |
| tcaatgttgtccttagcagc | nanog-pmc-mrna_39 | 30 nmol | Desalted | T(CAL Fluor Red | 7232.96 |
| tatctatagccagagacggc | nanog-pmc-mrna_40 | 30 nmol | Desalted | T(CAL Fluor Red | 7275.99 |
| tgaggttagattctaaaccc | nanog-pmc-mrna_41 | 30 nmol | Desalted | T(CAL Fluor Red | 7266 |
| agtatgttacagcttaaccc | nanog-pmc-mrna_42 | 30 nmol | Desalted | T(CAL Fluor Red | 7225.97 |
| caaaacagagccaaaaacgg | nanog-pmc-mrna_43 | 30 nmol | Desalted | T(CAL Fluor Red | 7296.03 |
| acaaccaacaaattagggga | nanog-pmc-mrna_44 | 30 nmol | Desalted | T(CAL Fluor Red | 7302.04 |
| gcagcaaatacaagacctct | nanog-pmc-mrna_45 | 30 nmol | Desalted | T(CAL Fluor Red | 7228.97 |
| gtagtactcatgtcattacg | nanog-pmc-mrna_46 | 30 nmol | Desalted | T(CAL Fluor Red | 7256.99 |
| ctcggtgaaatcagggtaaa | nanog-pmc-mrna_47 | 30 nmol | Desalted | T(CAL Fluor Red | 7340.05 |
| agctgtatatttactcatcg | nanog-pmc-mrna_48 | 30 nmol | Desalted | T(CAL Fluor Red | 7231.98 |

| Sequence | Sequence Name | Scale | Purification | Three Modificatic | MW Backbone |
|---|---|---|---|---|---|
| acagacaccaatggttggag | cdx2-pmc-mrna_1 | 30 nmol | Desalted | T(CAL Fluor Red | 7325.03 |
| acaagactctattagtaatg | cdx2-pmc-mrna_2 | 30 nmol | Desalted | T(CAL Fluor Red | 7274.03 |
| ccttccgtgattaacgagtg | cdx2-pmc-mrna_3 | 30 nmol | Desalted | T(CAL Fluor Red | 7257.97 |
| ggtccgggagcagacctcac | cdx2-pmc-mrna_4 | 30 nmol | Desalted | T(CAL Fluor Red | 7277.95 |
| gaggtagctcacgtacatgg | cdx2-pmc-mrna_5 | 30 nmol | Desalted | T(CAL Fluor Red | 7332.02 |
| ctgacgaagttctgcggcgc | cdx2-pmc-mrna_6 | 30 nmol | Desalted | T(CAL Fluor Red | 7283.96 |
| acgtggtaaccgccgtagtc | cdx2-pmc-mrna_7 | 30 nmol | Desalted | T(CAL Fluor Red | 7267.96 |
| gactgcgcgctgtccaagtt | cdx2-pmc-mrna_8 | 30 nmol | Desalted | T(CAL Fluor Red | 7258.95 |
| atacgctgccggccaggatg | cdx2-pmc-mrna_9 | 30 nmol | Desalted | T(CAL Fluor Red | 7292.97 |
| ctgctgtagcccatggctgc | cdx2-pmc-mrna_10 | 30 nmol | Desalted | T(CAL Fluor Red | 7234.92 |
| cgtttgcagcagcccagaag | cdx2-pmc-mrna_11 | 30 nmol | Desalted | T(CAL Fluor Red | 7276.97 |
| gttttcacttggctgccgag | cdx2-pmc-mrna_12 | 30 nmol | Desalted | T(CAL Fluor Red | 7264.96 |
| tcgatatttgtctttcgtcc | cdx2-pmc-mrna_13 | 30 nmol | Desalted | T(CAL Fluor Red | 7189.93 |
| gctggtggtccgtgtacacc | cdx2-pmc-mrna_14 | 30 nmol | Desalted | T(CAL Fluor Red | 7274.95 |
| ctttcctccggatggtgatg | cdx2-pmc-mrna_15 | 30 nmol | Desalted | T(CAL Fluor Red | 7264.96 |
| tttaacctgcctctcagaga | cdx2-pmc-mrna_16 | 30 nmol | Desalted | T(CAL Fluor Red | 7201.94 |
| tttgctctgcggttctgaaa | cdx2-pmc-mrna_17 | 30 nmol | Desalted | T(CAL Fluor Red | 7263.98 |
| ctctgggacacttctcagag | cdx2-pmc-mrna_18 | 30 nmol | Desalted | T(CAL Fluor Red | 7242.95 |
| gggaagacaccggactcaag | cdx2-pmc-mrna_19 | 30 nmol | Desalted | T(CAL Fluor Red | 7335.02 |
| tgccgctgcagaacccggtg | cdx2-pmc-mrna_20 | 30 nmol | Desalted | T(CAL Fluor Red | 7268.94 |
| catggctcagcctggaattg | cdx2-pmc-mrna_21 | 30 nmol | Desalted | T(CAL Fluor Red | 7282.98 |
| ctgaggagtctagcagagtc | cdx2-pmc-mrna_22 | 30 nmol | Desalted | T(CAL Fluor Red | 7332.02 |
| ccaggtctgtaggtctatgg | cdx2-pmc-mrna_23 | 30 nmol | Desalted | T(CAL Fluor Red | 7314 |
| gctcccatttttcctctgag | cdx2-pmc-mrna_24 | 30 nmol | Desalted | T(CAL Fluor Red | 7159.89 |
| atatttcttgaggccccaaa | cdx2-pmc-mrna_25 | 30 nmol | Desalted | T(CAL Fluor Red | 7225.97 |
| tggcagaaaaagccagatgg | cdx2-pmc-mrna_26 | 30 nmol | Desalted | T(CAL Fluor Red | 7374.07 |
| cggaggctttctgtctcctc | cdx2-pmc-mrna_27 | 30 nmol | Desalted | T(CAL Fluor Red | 7200.9 |
| ttctgccagtccggaatgaa | cdx2-pmc-mrna_28 | 30 nmol | Desalted | T(CAL Fluor Red | 7266.98 |
| gtgtggtcagtccaggcaat | cdx2-pmc-mrna_29 | 30 nmol | Desalted | T(CAL Fluor Red | 7323.01 |
| cctgcagatctaggaagaga | cdx2-pmc-mrna_30 | 30 nmol | Desalted | T(CAL Fluor Red | 7325.03 |
| ctcggctctagccagaggtg | cdx2-pmc-mrna_31 | 30 nmol | Desalted | T(CAL Fluor Red | 7283.96 |
| ctggacctccgagggccatg | cdx2-pmc-mrna_32 | 30 nmol | Desalted | T(CAL Fluor Red | 7268.94 |
| cttccgcagtgtaaaccttg | cdx2-pmc-mrna_33 | 30 nmol | Desalted | T(CAL Fluor Red | 7217.94 |
| agctttctatcttagctgcc | cdx2-pmc-mrna_34 | 30 nmol | Desalted | T(CAL Fluor Red | 7183.92 |
| gttctgcagtctttggtcag | cdx2-pmc-mrna_35 | 30 nmol | Desalted | T(CAL Fluor Red | 7279.98 |
| ctgtgcatacaccagccaag | cdx2-pmc-mrna_36 | 30 nmol | Desalted | T(CAL Fluor Red | 7220.94 |
| caggcctggagtccaataac | cdx2-pmc-mrna_37 | 30 nmol | Desalted | T(CAL Fluor Red | 7260.97 |
| ctaaacaagtccctgttcgg | cdx2-pmc-mrna_38 | 30 nmol | Desalted | T(CAL Fluor Red | 7226.95 |
| gacatacattcagcccagag | cdx2-pmc-mrna_39 | 30 nmol | Desalted | T(CAL Fluor Red | 7244.97 |
| aggaagtccaggttggctct | cdx2-pmc-mrna_40 | 30 nmol | Desalted | T(CAL Fluor Red | 7323.01 |
| ctctggcttggatgttacac | cdx2-pmc-mrna_41 | 30 nmol | Desalted | T(CAL Fluor Red | 7248.96 |
| catggatccagaaggcttta | cdx2-pmc-mrna_42 | 30 nmol | Desalted | T(CAL Fluor Red | 7291.01 |
| tctgaaatctggaaagctca | cdx2-pmc-mrna_43 | 30 nmol | Desalted | T(CAL Fluor Red | 7275.01 |
| gcagacaaaatctgaagata | cdx2-pmc-mrna_44 | 30 nmol | Desalted | T(CAL Fluor Red | 7317.06 |
